# Modeling Spatio-temporal Dynamics of Chromatin Marks

**DOI:** 10.1101/239442

**Authors:** Petko Fiziev, Jason Ernst

## Abstract

To model spatial changes of chromatin mark peaks over time we developed and applied ChromTime, a computational method that predicts regions for which peaks either expand or contract significantly or hold steady between time points. Predicted expanding and contracting peaks can mark regulatory regions associated with transcription factor binding and gene expression changes. Spatial dynamics of peaks provided information about gene expression changes beyond localized signal density changes. ChromTime detected asymmetric expansions and contractions, which for some marks associated with the direction of transcription. ChromTime facilitates the analysis of time course chromatin data in a range of biological systems.

## BACKGROUND

Genome-wide mapping of histone modifications (HMs) and related chromatin marks using chromatin immunoprecipitation coupled with high-throughput sequencing (ChIP-seq) and DNA accessibility through assays for DNase I hypersensitivity[1] (DNase-seq) or transposase-accessible chromatin[2] (ATAC-seq) assays have emerged as a powerful approach to annotate genomes and study cell states[3–5]. Through the efforts of large consortia projects such as the ENCODE[6], Roadmap Epigenomics[7] and BLUEPRINT[8] as well as individual labs[9–11] multiple different chromatin marks have been mapped across more than a hundred different cell and tissue types. These maps have yielded numerous insights into gene regulation and genetic and epigenetic association with disease[12–16].

While many mapping efforts have largely focused on single or unrelated cell and tissue types[3, 6], a growing number of biological processes have been studied with temporal epigenomic data using assays such as ChIP-seq, ATAC-seq or DNase-seq in a time course that map chromatin marks at consecutive stages during particular biological processes. Such datasets have been generated for a wide range of biological settings including T cell development[17], adipogenesis[18], hematopoiesis[19, 20], macrophage differentiation[21], neural differentiation[12], cardiac development[22, 23], somatic cell reprogramming[24–27], embryogenesis[28] and many others[7, 29–37]. The output of these experiments presents a unique opportunity to study the spatio-temporal changes of epigenetic peaks and associated regulatory elements. However, almost all computational methods designed or applied to epigenomic data have been developed based on single or multiple unrelated samples. For example, continuous regions of enrichments of single marks are detected by peak or domain calling methods[38–42]. In cases when multiple chromatin marks are mapped in the same cell type, methods such as ChromHMM[43] and Segway[44] can be used to produce genome-wide chromatin state annotations. In addition, methods have been developed for pairwise comparisons of ChIP-seq signal data by differential peak calling[45, 46].

In the context of time course chromatin data, only a few methods have been proposed that consider temporal dependencies between samples. One such method, TreeHMM[47], produces a chromatin state genome annotation similar to ChromHMM and Segway, while taking into account a tree-like structure that captures lineage relationships between the input cell types in order to potentially derive a more consistent annotation across samples. Another method, GATE[30], produces a genome annotation based on clustering fixed length genomic loci that can be modeled with the same switch from one chromatin state to another over time.

One important limitation of methods for pairwise comparison or time course modeling of chromatin data is that they do not directly consider or model spatial changes in the genomic territory occupied by chromatin marks over time. Spatial properties of genomic peaks continuously marked by HMs have gained increasing attention as a potentially important characteristic of chromatin marks. For example, long peaks of H3K27ac have been associated with active cell type specific locus control regions termed super-enhancers or stretch enhancers in a number of cell types[48, 49]. Also, the length of H3K4me3 peaks has been associated with transcriptional elongation and consistency of cell identity genes[50]. In the context of cancer, long H3K4me3 peaks have been linked to transcriptional elongation and enhancer activity at tumor suppressor genes and have been observed to be significantly shortened in tumor cells[51]. Long H3K4me3 domains have been implicated to mark loci involved in psychiatric disorders[52]. Expanded domains of H3K27me3 and H3K9me3 marks have been shown to be characteristic of terminally differentiated cells compared to stem cells[53]. These studies suggest that length of epigenetic peaks is a dynamic feature that can correlate with activity of putative functional elements regulating specific genes. Computational methods that do not explicitly reason about the spatial changes of chromatin marks have significant limitations for studying the dynamics of these properties because they are unable to detect territorial changes that might be associated with redistribution of signal or identify asymmetric directional peak boundary movements.

In this work, we present ChromTime, a novel computational method for detection of expanding, contracting and steady peaks, which can detect patterns of changes in the genomic territory occupied by chromatin mark peaks from time course sequencing data (**Fig 1A**). We applied ChromTime to a diverse set of data from different developmental, differentiation and reprogramming time courses. Predicted expansions and contractions in general mark regulatory regions associated with changes in transcription factor (TF) binding or gene expression. ChromTime enables studying the directionality of spatial dynamics of chromatin mark peaks relative to other genomic features, which existing computational approaches do not directly address. Our results show that the direction of predicted expansions and contractions correlates with direction of transcription near transcription start sites (TSSs). ChromTime is a general method that can be used to analyze time course chromatin data from high-throughput sequencing assays such as from ChIP-seq, ATAC-seq and DNase-seq for a wide range of biological systems to gain insights into the dynamics of gene regulation.

**Figure 1:**
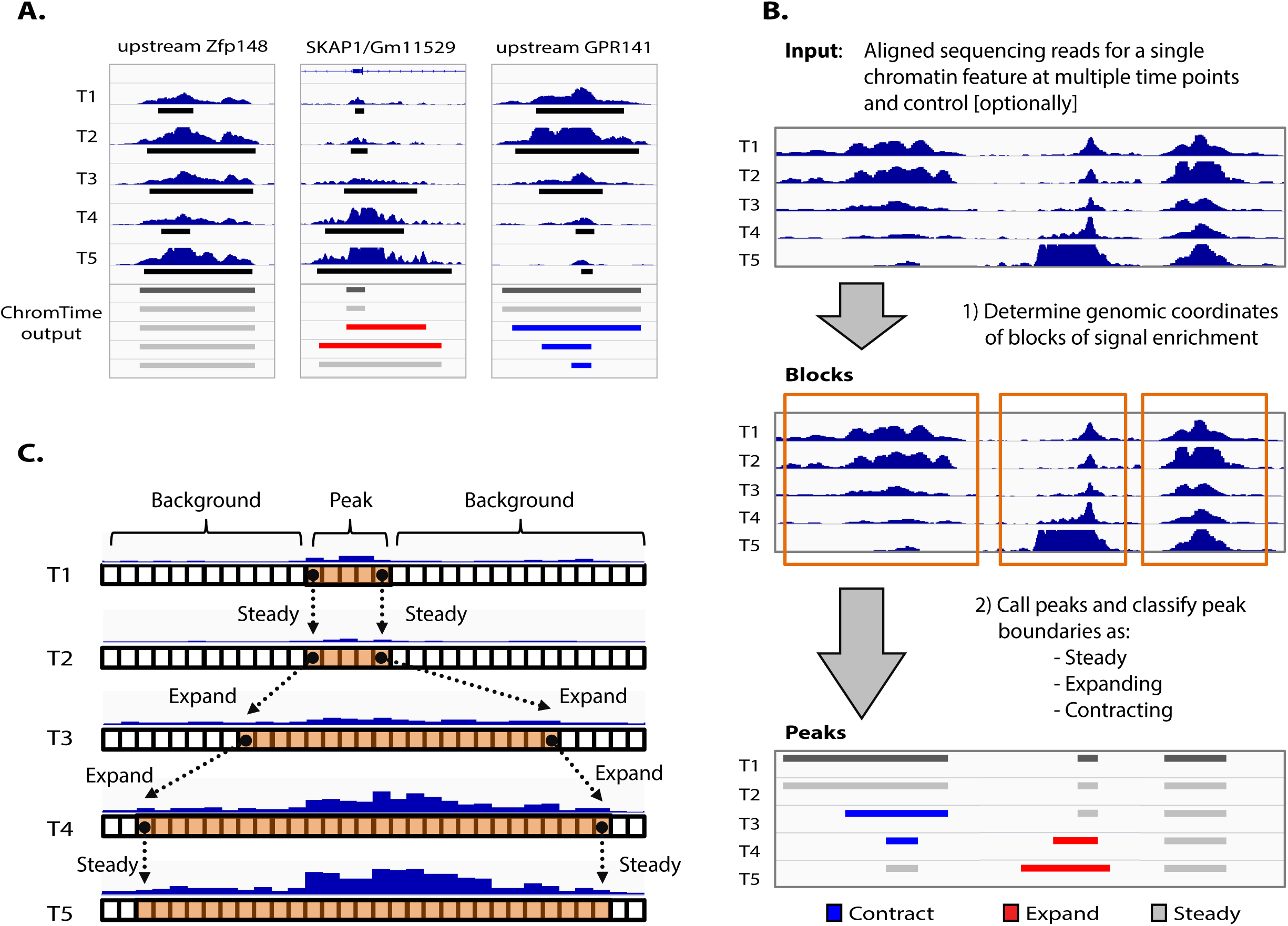
Overview of the ChromTime method. **(A)** Examples of H3K4me2 peaks with steady, expanding and contracting boundary dynamics, shown from left to right respectively, across five time points during mouse T cell development[17]. Time points 1, 2 and 3 correspond to in vitro differentiated T cell precursors (FLDN1, FLDN2a and FLDN2b), whereas time points 4 and 5 correspond to in vivo purified thymocytes (ThyDN3 and ThyDP). Normalized ChIP-seq signal, MACS2[38] peaks (black rectangles) and ChromTime output are shown for each time point. Peaks upstream of Zfp148 gene are called steady by ChromTime despite fluctuations of MACS2 peak boundaries. In contrast, ChromTime calls a peak at the Skap1/GM11529 promoter to expand after time points 2 and 3. Conversely, ChromTime calls a peak upstream of GPR141 gene to contract after time points 2, 3, and 4. **(B)** Overview of the ChromTime method. During the block finding stage, input foreground and, optionally, control reads are used to determine blocks of signal enrichment. In the dynamics prediction stage, for each block, peak boundary positions are predicted at each time point and peak boundary dynamics are predicted at each pair of consecutive time points. **(C)** Schematic of predicting dynamics for one block. Boxes represent genomic bins at each time point. Foreground signal is depicted as blue bars for each bin whose height represents the number of reads mapped to the bin. ChromTime learns a probabilistic mixture model from the input data to partition each block at each time point into peak and background components. Bins in the peak component (orange) mark peaks of signal enrichment whereas those in the background component (white) mark flanking background signal. The movement of the boundaries on the left and the right side of peaks between consecutive time points are estimated by reasoning jointly about the input data from all time points.

## RESULTS

### Model for detecting expanding, contracting and steady peaks from temporal chromatin data

We developed a computational method, ChromTime (https://github.com/ernstlab/ChromTime), designed for systematic detection of expansions, contractions and steady peaks from time course chromatin data for single chromatin features (**Methods, Fig 1B**). ChromTime takes as input a set of genomic coordinates of aligned sequencing reads from foreground and, optionally, control experiments over the time course. The foreground experiments are data from a chromatin sequencing assay such as ChIP-seq, ATAC-seq or DNase-seq performed at a series of time points. The method consists of two stages – block finding and dynamics prediction. During the block finding stage, ChromTime determines continuous genomic regions (blocks) that may contain peaks of foreground signal enrichment during the time course (**Fig S1A-B**). To achieve this, ChromTime partitions the genome into fixed length bins and counts the number of foreground and control reads that map to each bin at each time point. Nearby bins that show significant enrichment are joined into continuous intervals, which subsequently are grouped into blocks if they overlap across time points. As a result, large portions of the genome that are likely to contain background noise at all time points are filtered out, so that peak boundary dynamics are determined within a subset of the genome potentially enriched for the chromatin mark.

During the dynamics prediction stage, for each block, ChromTime determines the most likely positions of the peak boundaries at each time point and whether the peak expands, contracts or holds steady at each boundary between consecutive time points. The method uses a probabilistic mixture model to partition the signal within each block at each time point into background and peak components (**Fig 1C, S1C**) by reasoning jointly about the data from all time points in the time course. The method assumes that central positions in blocks are more likely to be enriched for foreground reads and thus the peak component is flanked by the background components (**Fig S1D**). The number of sequencing reads in bins from each component at each time point is modelled with different negative binomial distributions that can account for the local abundance of control reads. Furthermore, between any two consecutive time points the boundaries of the peaks are assumed to follow one of three possible dynamics: steady, expand or contract. For steady dynamics, the peak boundaries are enforced to have the same genomic position. For expanding and contracting dynamics, the number of genomic bins that the peak boundaries move between the two time points is modelled with different negative binomial distributions which depend on the pair of time points and the corresponding dynamic. ChromTime models time points that have no bins in the peak component with zero length peaks. Thus, appearances of peaks, except at the first time point, are modeled as expansions from zero length peaks and the disappearances of peaks are modeled as contractions to zero length peaks. Each dynamic is also assumed to have a prior probability which captures information about its genome-wide frequency at each time point.

All model parameters are learned jointly from the whole time course. As a result, ChromTime can adapt to different boundary movements, dynamics frequencies and noise levels across experiments and biological systems. The estimated parameters are used to make a prediction for each block for the most likely positions of the peak boundaries and the corresponding boundary dynamics that had generated the signal within the block. The final output contains predicted peak boundaries annotated and colored by their assigned dynamics, which can be used for downstream analysis with existing tools and visualized in genome browsers (**Fig 2, S2,** https://github.com/ernstlab/ChromTime).

**Figure 2:**
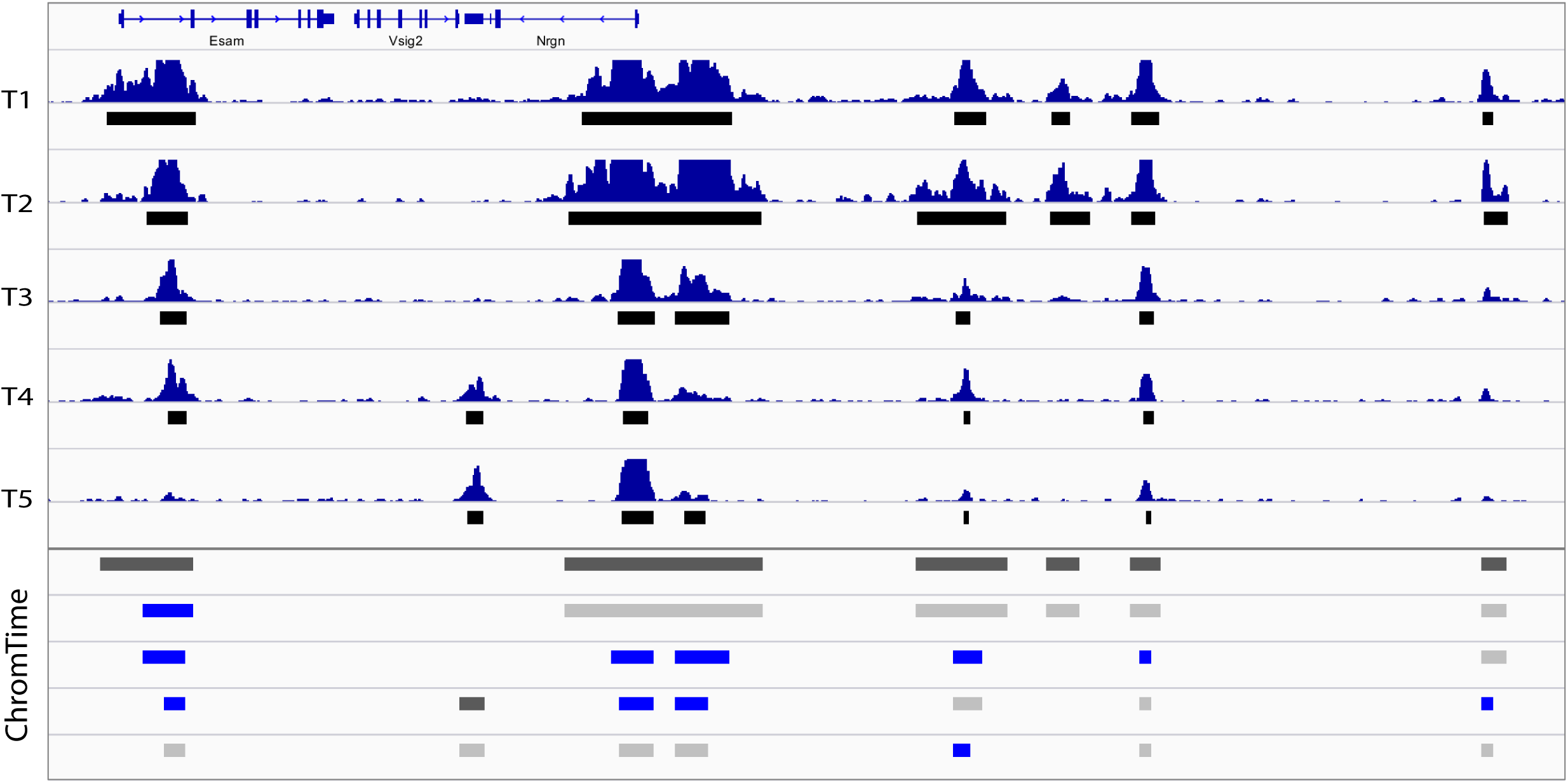
Sample output from ChromTime with contracting peaks. Genome browser screenshot with sample output of ChromTime for H3K4me2 from the T cell development time course in mouse[17] with 5 time points at the Esam/Vsig2/Nrgn locus. Time points 1, 2 and 3 correspond to in vitro differentiated T cell precursors (FLDN1, FLDN2a, and FLDN2b), whereas time points 4 and 5 correspond to in vivo purified thymocytes (ThyDN3 and ThyDP). The input ChIP-seq signal and MACS2[38] peaks (black boxes under each signal track) are shown in the upper panel of the screenshot. The ChromTime predicted peaks colored by their boundary dynamics for each block at each time point are shown in the bottom panel. The first peak in each block is colored in dark grey. Each subsequent peak is colored with respect to the predicted dynamic relative to its previous time point. Peaks with steady boundaries on both sides are shown in light grey, and those with at least one contracting boundary are shown in blue. Nearby peaks that touch boundaries are visualized as one peak by the genome browser. Not shown in the figure are expanding peaks, peaks at single time points and peaks with opposite dynamics (EXPAND on the left and CONTRACT on the right, or vice versa), which would be colored in red, orange and black, respectively. See **Fig S2** for examples of predicted expanding peaks.

### Reproducibility of ChromTime predictions and association with changes in gene expression, TF binding and DNaseI hypersensitivity sites

To investigate the reproducibility of ChromTime predictions, we applied ChromTime separately to two biological replicate datasets for the H3K4me2 and the H3K(9/14)ac marks in T cell development in mouse[17] and confirmed on average strong enrichment for the same ChromTime annotations co-localizing across replicates (**Fig S3**). We then applied the method to data from pooled replicates for the H3K4me2 mark from the mouse T cell development study[17], to data for the H3K4me3 and H3K27ac marks from a study on stem cell reprogramming in human[24], to ATAC-seq data from a mouse stem cell reprogramming time course[27] and to a human fetal brain development time course that we constructed from DNase-seq datasets from Roadmap Epigenomics[7]. To investigate the biological relevance of ChromTime predictions, for blocks overlapping TSSs we examined changes in the corresponding gene expression. Peaks with predicted expanding and contracting boundaries that overlap annotated TSSs associated with increase and decrease, respectively, in gene expression (**Fig 3, S4**). Additionally for all chromatin marks, we examined enrichments of TF binding sites across all blocks[6, 17, 27], and in the case of HMs, also enrichments of DNaseI hypersensitivity sites (DHSs)[7]. Predicted peaks with expanding and contracting boundaries enriched for sites bound by important regulators in each biological system as well as sites bound by generic TFs in a cell type specific manner. Expanding and contracting HM peaks also enriched for cell type specific DHSs. Furthermore, peaks with predicted steady boundaries showed enrichment for TF binding sites that are shared between the first and the last time point in the corresponding time courses, which mark potentially stable regulatory elements. Similar enrichments in the case of HM peaks were also seen for shared DHSs.

**Figure 3:**
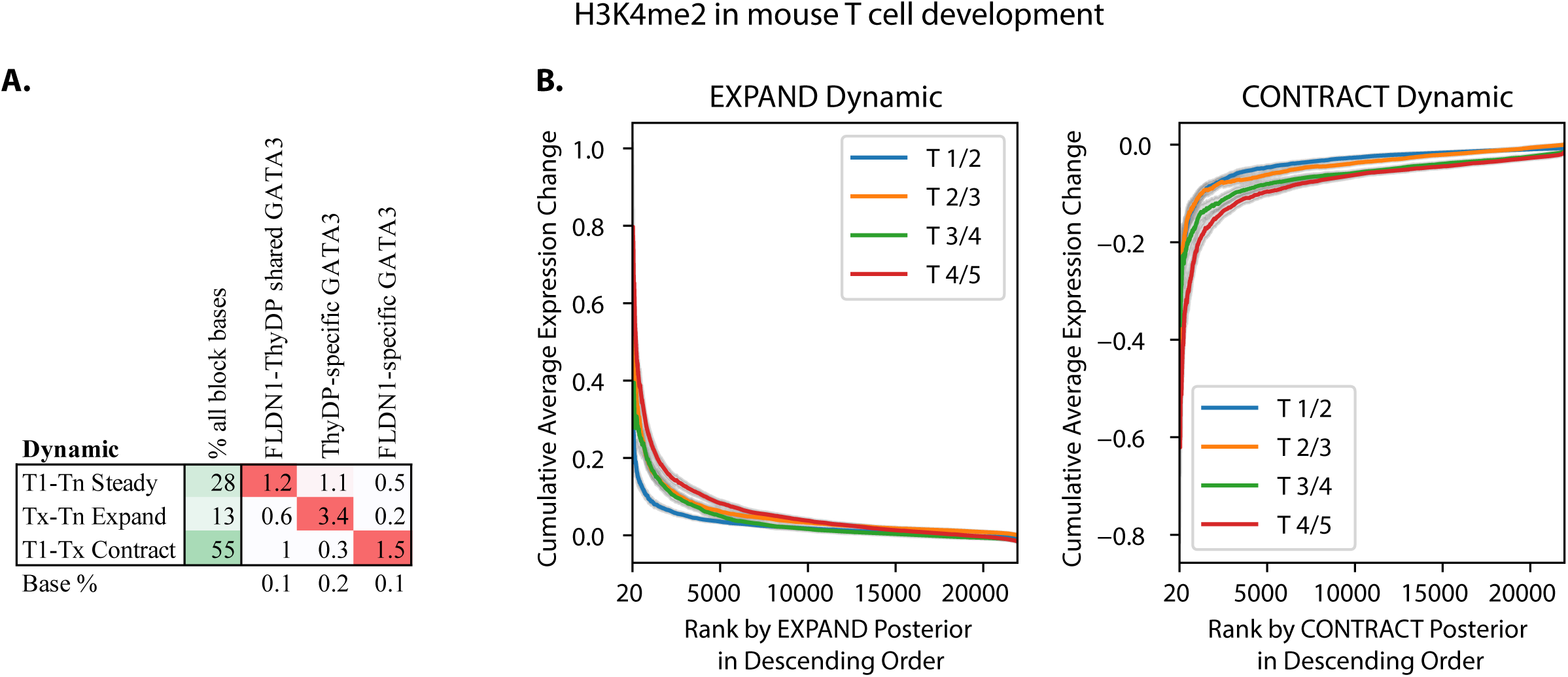
Changes in GATA3 binding and gene expression at predicted H3K4me2 dynamics in T cell development. (**A**) Fold enrichments of cell type specific and shared peaks of GATA3, which is a master regulator in T cell development[17], are shown for three sets of blocks with predicted H3K4me2 peaks: 1) blocks with peaks present at all time points whose boundaries hold steady on both sides throughout the whole time course (T1-Tn Steady); 2) blocks with non-contracting peaks whose boundaries expand between at least one pair of consecutive time points and have a peak at the last time point (Tx-Tn Expand); and 3) blocks with non-expanding peaks whose boundaries contract between at least one pair of consecutive time points and have a peak at the first time point (T1-Tx Contract). First column shows the percentage of bases out of all bases covered by peaks of the set. Last row shows the baseline percentage for each feature out of all bases covered by ChromTime peaks at any time point. Percentages are colored from 0 (white) to 100 (green). Fold enrichments in each column are colored from 1 (white) to the maximum value in the column (red). FLDN1 and ThyDP denote differentiated T cell precursors and purified thymocytes, which are the first and the last time point, respectively. **(B)** Boundaries of predicted H3K4me2 peaks in blocks with at least one predicted non-zero length peak overlapping annotated TSSs were sorted in decreasing order by their posterior probability for EXPAND dynamic (left plots) and CONTRACT dynamic (right plots) at each pair of consecutive time points (**Supplementary Methods**). Gene expression differences between consecutive time points were calculated as the average difference across all genes with overlapping TSSs for each block. For each posterior rank (X-axis) the plot shows the cumulative average gene expression difference (Y-axis). Expanding boundaries associated with increase of gene expression and contracting boundaries associated with decrease of gene expression. Shaded regions correspond to 95% confidence intervals.

### Predicted spatial dynamics by ChromTime associate better with gene expression changes compared to boundary position changes of peaks called from individual time points in isolation

We next investigated whether ChromTime’s approach for reasoning jointly about the whole time course increases power to detect associations with gene expression compared to considering boundary differences of peaks at consecutive time points called in isolation. Specifically, we analyzed gene expression changes of genes with TSSs overlapping ChromTime peaks in relation to posterior probabilities for expansions and contractions compared to boundary differences of peaks called with ChromTime from data from individual time points in isolation. We investigated this in the context of H3K4me2 peaks in mouse T cell development[17] and for H3K4me3 peaks in stem cell reprogramming in human[24]. In most cases, ranking boundary changes of peaks in blocks with at least one non-zero length peak by their predicted ChromTime posterior probabilities for expansions and contractions associated on average with larger gene expression changes compared to ranking boundaries directly based on the change in the genomic positions of the boundaries of ChromTime peaks called at individual time points in isolation (**Fig 4, S5A**). These results also held when using peaks from two different peak callers, MACS2[38] and SICER[40] applied on data from individual time points (**Fig S5B-C**).

**Figure 4:**
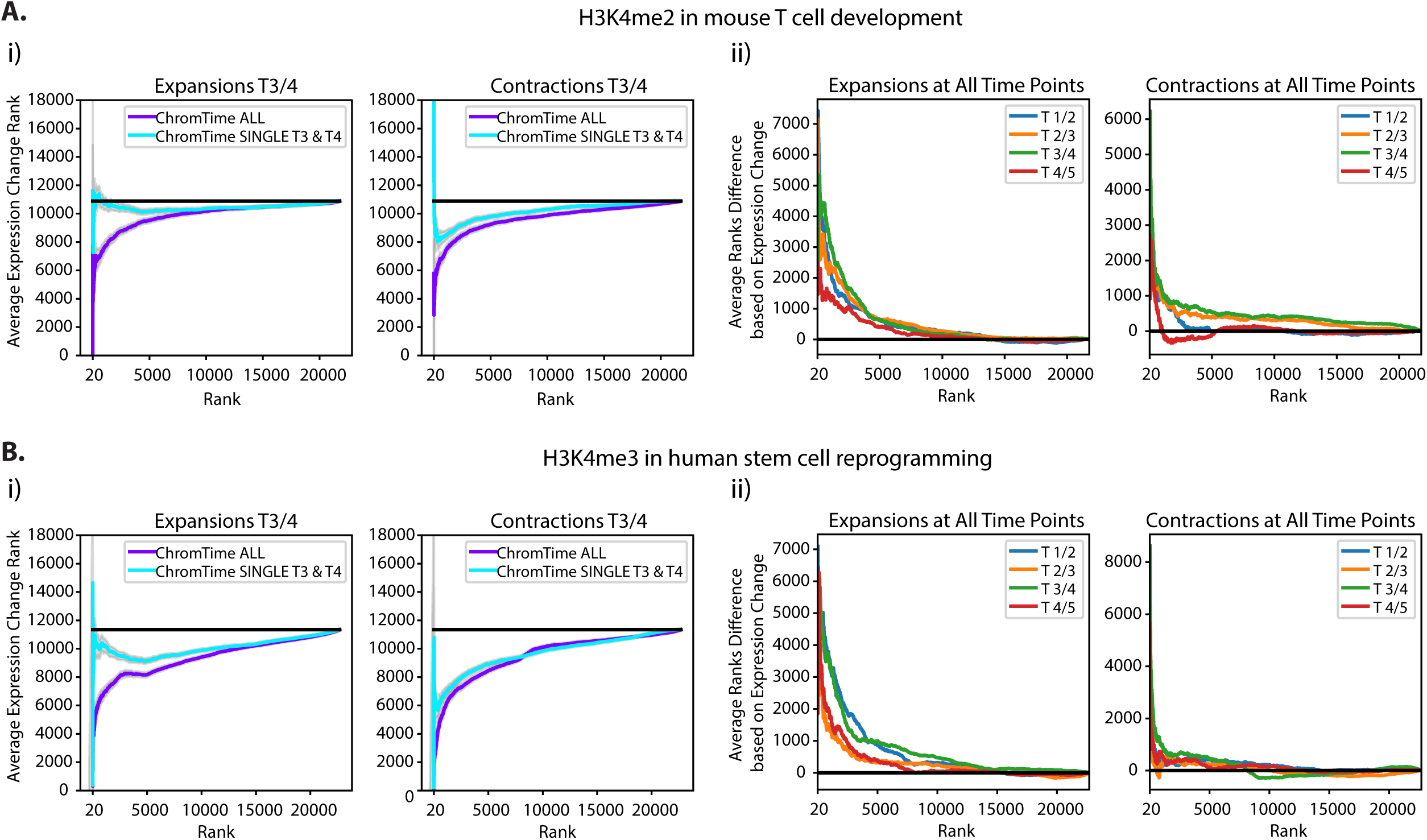
ChromTime predictions associate better with expression changes than boundary movements of peaks called in isolation. **(A)** For H3K4me2 in mouse T cell development[17] ChromTime was applied once with data from all time points (ChromTime ALL), and once with single time points in isolation (ChromTime SINGLE; **Supplementary Methods**). Time points 1, 2 and 3 correspond to T cell precursors, whereas 4 and 5 to purified thymocytes. Peaks called by both procedures overlapping annotated TSSs were analyzed for their relationship with gene expression changes. (**i**) (Left) Comparison of agreement with expression for expansions when applying ChromTime ALL and ChromTime SINGLE for the change between time points 3 and 4. Peak boundaries were sorted in decreasing order of their EXPAND posterior probabilities from ChromTime ALL and compared to sorting them in decreasing order of the difference of peak boundary positions in ChromTime SINGLE peaks with positive differences in boundary positions indicating peaks expanding with time. Each boundary was also ranked by the average gene expression difference of TSSs overlapping its block in decreasing order with positive expression differences indicating gain with time. The cumulative average boundary rank of expression change (Y-axis) is shown for the boundary change ranking for ChromTime ALL and ChromTime SINGLE (X-axis). Low Y-values indicate stronger association with expression changes. Black line shows expected average expression change rank. Shaded regions indicate 95% confidence intervals. Plots for other time points can be found in **Fig S5**. (Right) Analogous to left plots for Contract posterior probabilities for ChromTime ALL, increasing order of the difference of boundary change positions for ChromTime SINGLE, and increasing order of expression changes. (**ii**) Differences between ChromTime ALL and ChromTime SINGLE values shown in (**i**) between time points 3 and 4 as well as for all other pairs of time points. Positive values correspond to boundary ranks for which ChromTime ALL posteriors better associate with gene expression changes than boundary movements of ChromTime SINGLE peaks. Black lines show expected difference of zero between random rankings. **(B)** As in (**A**) for H3K4me3 in human stem cell reprogramming[24]. Time points correspond to human inducible and immortalized fibroblasts-like (hiF-T) cells, hiF-T at 5, 10 and 20 days after induction, and human induced pluripotent stem cells (hIPSC).

### Spatial dynamics contain information about gene expression changes between consecutive time points not captured by corresponding pairwise signal density changes

We next investigated whether there is additional information in ChromTime predictions beyond what can be captured by pairwise signal density changes or by differential peak calls. For this analysis, we focused on H3K4me2 in mouse T cell development[17] and H3K4me3 in human stem cell reprogramming[24]. For pairs of consecutive time points, we computed the change in signal density in the region starting at the left most and ending at the right most predicted peak boundary in the block (**Supplementary Methods**). We associated the signal density changes with gene expression changes at the nearest TSS within 50kb of each block and computed the average gene expression change as a function of the signal density change within blocks (**Fig 5**). We found that locations with the same signal density change can associate with significantly different average gene expression changes of proximal genes depending on the predicted ChromTime dynamics. Notably, bidirectional expansions, expansions occurring on both sides of a peak, associated for a range of signal density changes with greater average increase in gene expression than unidirectional expansions, those expansions occurring on one side with steady on the other, when controlling for the signal density change. These unidirectional expansions in turn associated for a range of signal density changes with greater expression change than steady regions, those regions with a steady call on both sides of a peak, when controlling for the signal density change. We observed a similar relationship for contractions and decrease of gene expression. These results were replicated also after substituting ChIP-seq signal density changes with differential peak scores from two differential peak calling methods, SICER[40] and MACS2[38] (**Fig S6A-B**). Therefore, ChromTime predictions can provide additional information about gene expression changes beyond what is contained in the corresponding signal density changes as measured directly or by utilizing differential peak calling procedures.

**Figure 5:**
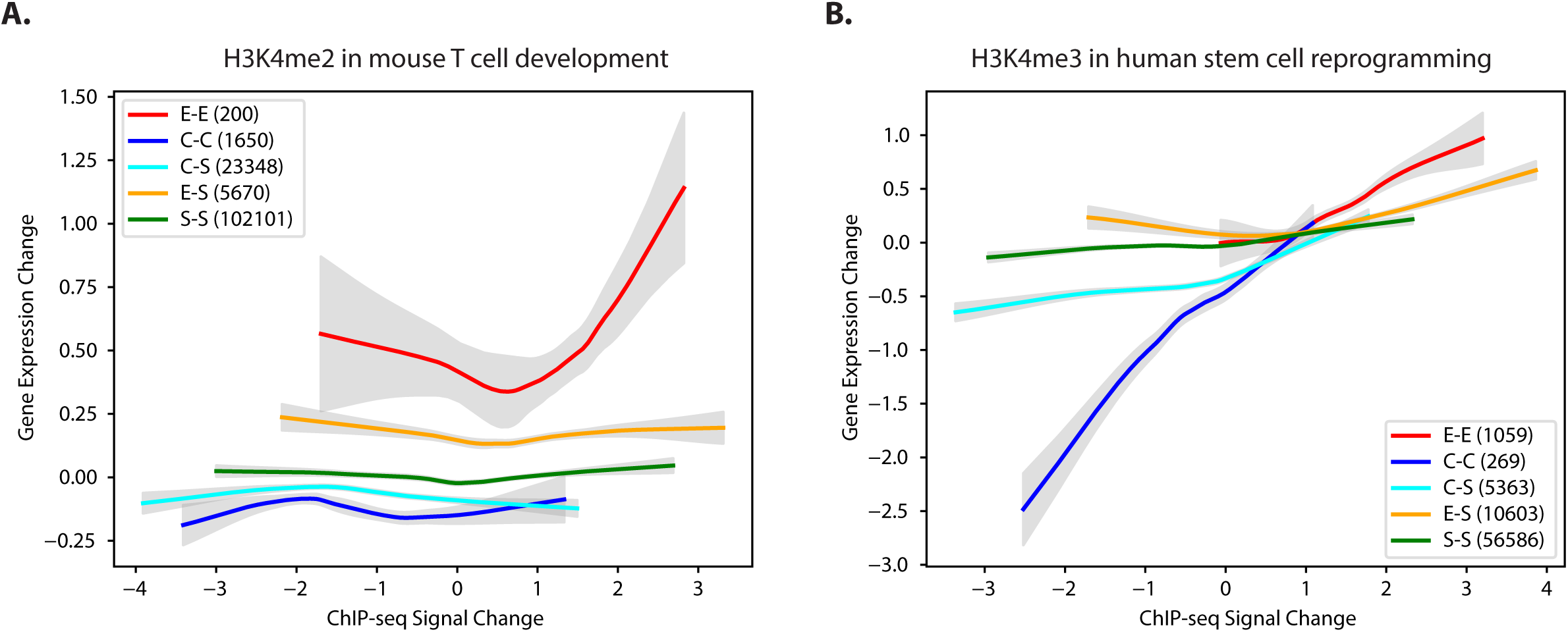
Spatial dynamics can contain additional information about gene expression changes beyond signal density changes. Gene expression change is plotted as function of ChIP-seq signal density change after loess smoothing for each predicted ChromTime dynamic for **(A)** H3K4me2 dynamics in T cell development in mouse[17]; and **(B)** in H3K4me3 dynamics in stem cell reprogramming in human[24] (**Supplementary Methods**). Peaks of each type of dynamics were pooled from all time points for this analysis. Peaks with asymmetric dynamics E/S and S/E were pooled together in the “E-S” group. Similarly C/S and S/C peaks were pooled in the “C-S” group. The total number of peaks in each group is shown in parenthesis. In both systems, for a range of signal density changes, peaks with the same signal density change associated with different gene expression changes depending on the predicted spatial dynamic. Shaded regions represent 95% confidence intervals.

### Spatial dynamics are correlated between multiple chromatin marks

Previous studies have shown that the locations of different chromatin marks can be correlated[3, 54]. In this context, we tested whether multiple chromatin marks can also exhibit jointly the same type of spatio-temporal dynamics. For this purpose, we compared the genomic locations of predicted expansions, contractions and steady peaks for different chromatin marks within the same time course. We focused on three previously published time courses – stem cell reprogramming in human[24], stem cell reprogramming in mouse[27] and adipogenesis in mouse[18], where multiple chromatin marks were mapped (**Fig 6, S7**). In all three datasets, we observed that predicted expansions co-localized preferentially for H3K4me2, H3K4me3 and H3K27ac and to a lesser extent for H3K4me1 and similarly for predicted contractions and steady peaks. In contrast, different predicted spatial dynamics for H3K36me3 and H3K27me3 tended to occupy distinct locations. In addition, in mouse reprogramming[27], ChromTime predicted dynamics of ATAC-seq, H3K4me2, H3K4me3, H3K27ac, H3K9ac and to a lesser extent of H3K4me1 and H3K79me2 peaks co-localized preferentially (**Fig S7**). These results suggest that spatial dynamics of chromatin marks are coordinated at least at a subset of genomic locations.

**Figure 6:**
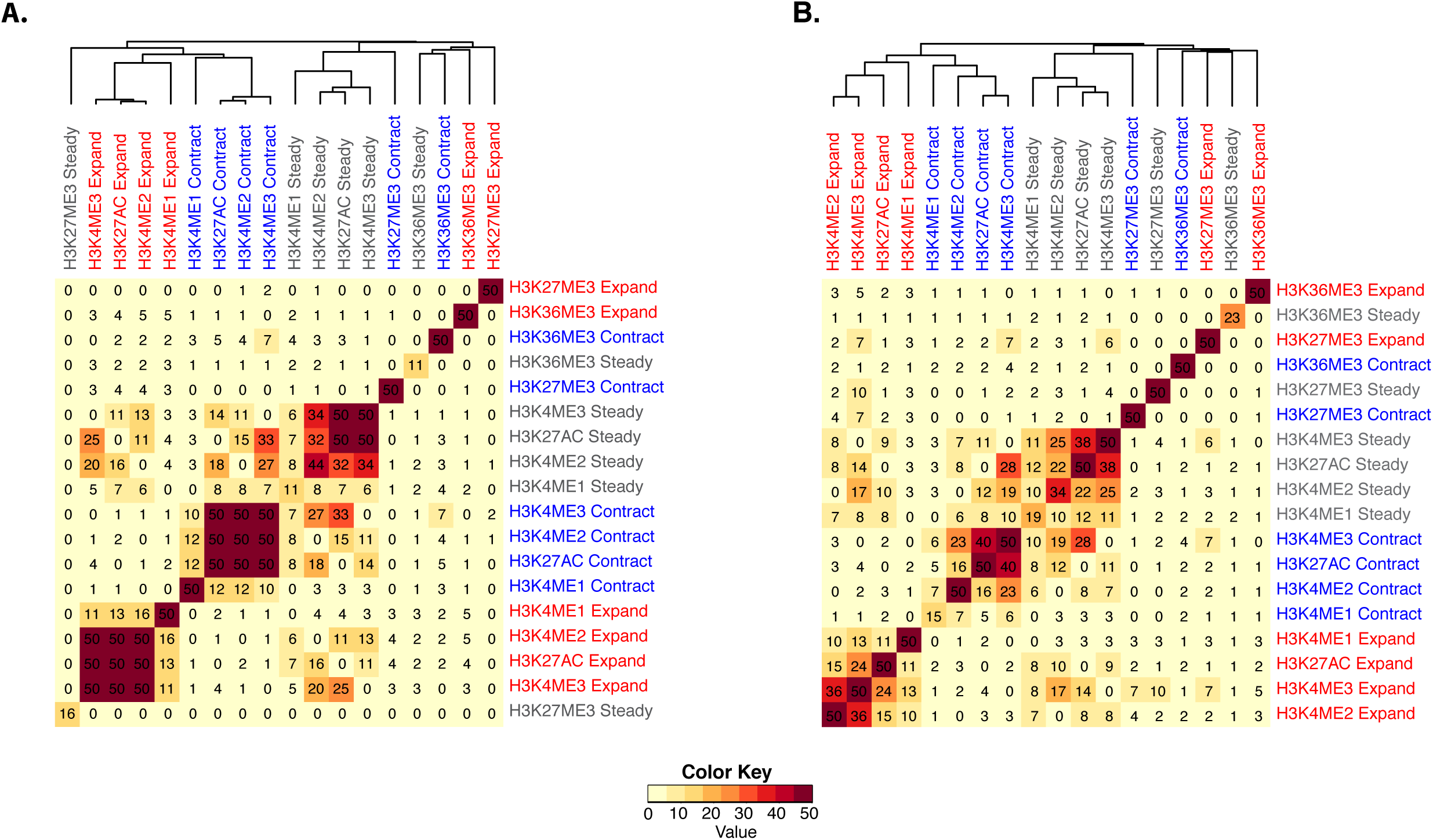
Spatial dynamics of multiple different chromatin marks co-localize within a time course. Hierarchical clustering with optimal leaf ordering[60] of the geometric average fold enrichments taken across all time points of the overlap of every pair of predicted spatial dynamics for mapped HMs in **(A)** mouse adipogenesis[18] and **(B)** human stem cell reprogramming[24]. At each pair of time points, “Expand” and “Contract” dynamics are defined as all peaks that are predicted as either unidirectional or bidirectional expansions and contractions, respectively, whereas “Steady” dynamics are defined as all peaks that have predicted steady boundaries at both sides. Peaks with “Expand” dynamic on one side and “Contract” dynamic on the other were excluded from this analysis. In both datasets, expansions, contractions and steady peaks of H3K4me2, H3K4me3 and H3K27ac and to a lesser extent of H3K4me1 tend to cluster together within each of the three classes, whereas spatial dynamics of H3K27me3 and H3K36me3 peaks tend to occupy different locations. All enrichments were capped at 50 before clustering.

### Direction of expansions and contractions is correlated with direction of transcription

ChromTime can predict unidirectional expansions and contractions, which enables analysis of directionality of spatial dynamics of peaks, an aspect of chromatin regulation that has not been previously systematically explored. To investigate this, we applied ChromTime on data from 13 previously published studies from a variety of developmental, differentiation and reprogramming processes (**Table 1**) for nine different HMs including narrow and broad marks, for Pol2, ATAC-seq and DNase-seq. We observed that unidirectional expansions and contractions are predicted in most cases on average to be the majority of all expansions and contractions, respectively, at a given pair of consecutive time points (**Fig S8**). One hypothesis for the prevalence of asymmetric boundary movements for the promoter associated chromatin marks is that the direction of boundary movements is associated with the asymmetry of transcription initiation in promoter regions. To test this hypothesis, for each dataset we compared the prevalence of each class of unidirectional dynamics as a function of its distance to the nearest annotated TSS and the orientation of the corresponding gene (**Fig 7**). Consistent with our hypothesis, for H3K4me3, H3K4me2, H3K(9/14)ac, H3K79me2, and for Pol2, we found that unidirectional expansions that expand into the gene body (i.e. in the same direction as transcription) were substantially more frequently found in proximity of TSSs compared to unidirectional expansions in the opposite direction. Moreover, this difference was not observed for expansions that are distal from TSSs. Similarly, in most cases for these data unidirectional contractions that contract towards the TSS of the nearest gene (i.e. in the opposite direction of transcription) were substantially more frequent compared to unidirectional contraction in the opposite direction in proximity of TSSs, whereas their frequencies at distal sites showed much smaller differences. HMs H3K27ac, H3K4me1 and H3K27me3, some ATAC-seq datasets and peaks in the DNase-seq time course exhibited the same trend, but to a smaller degree.

**Table 1:**
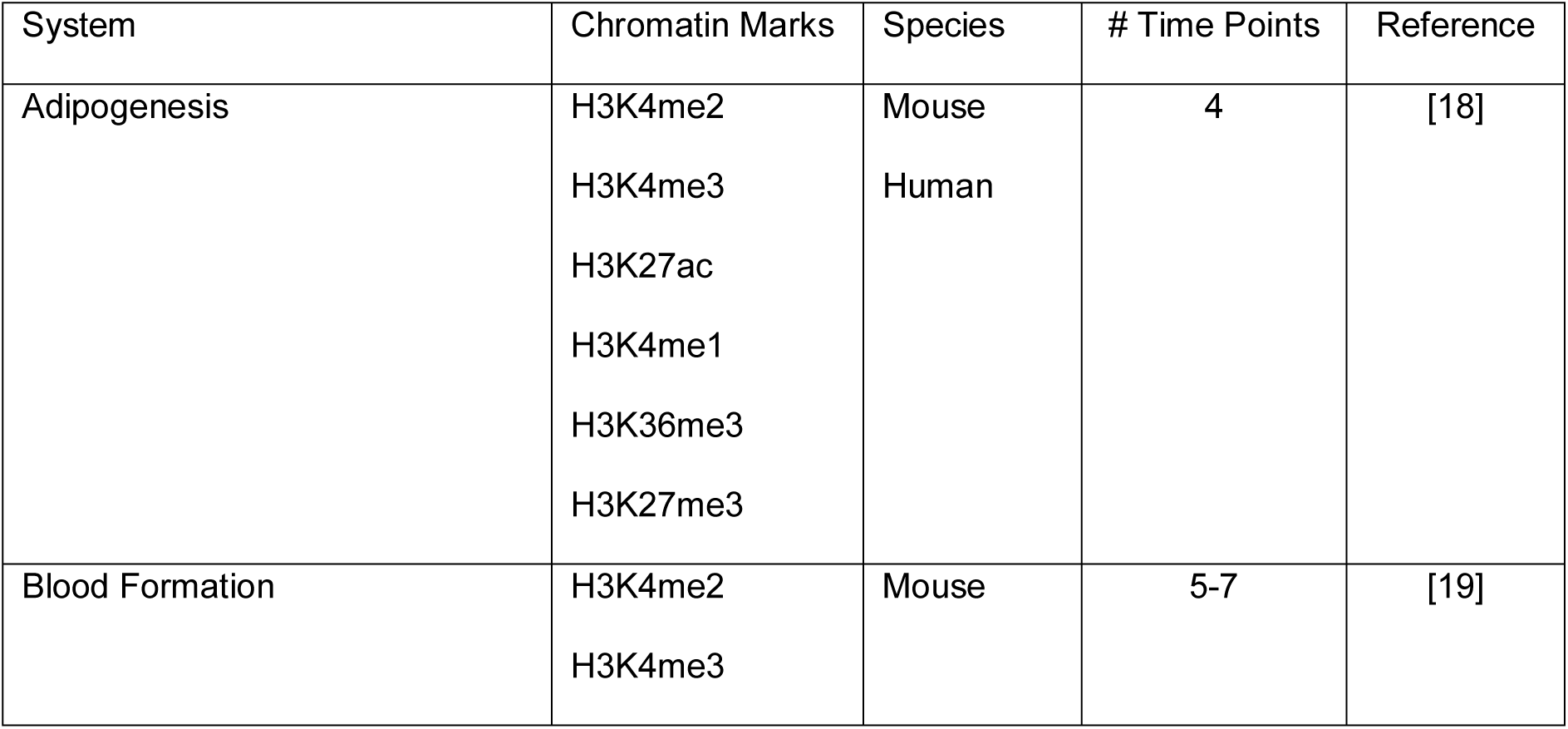
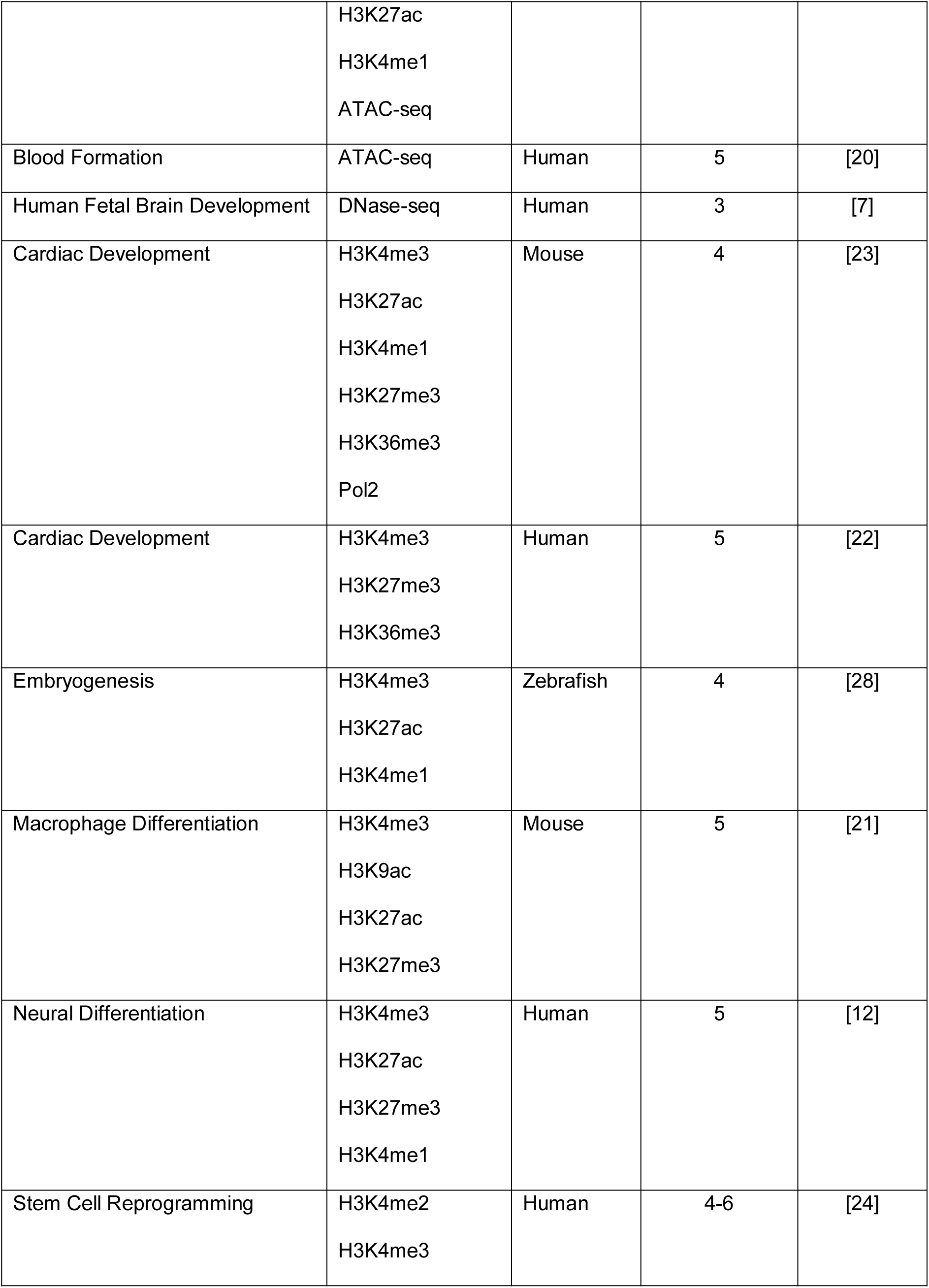
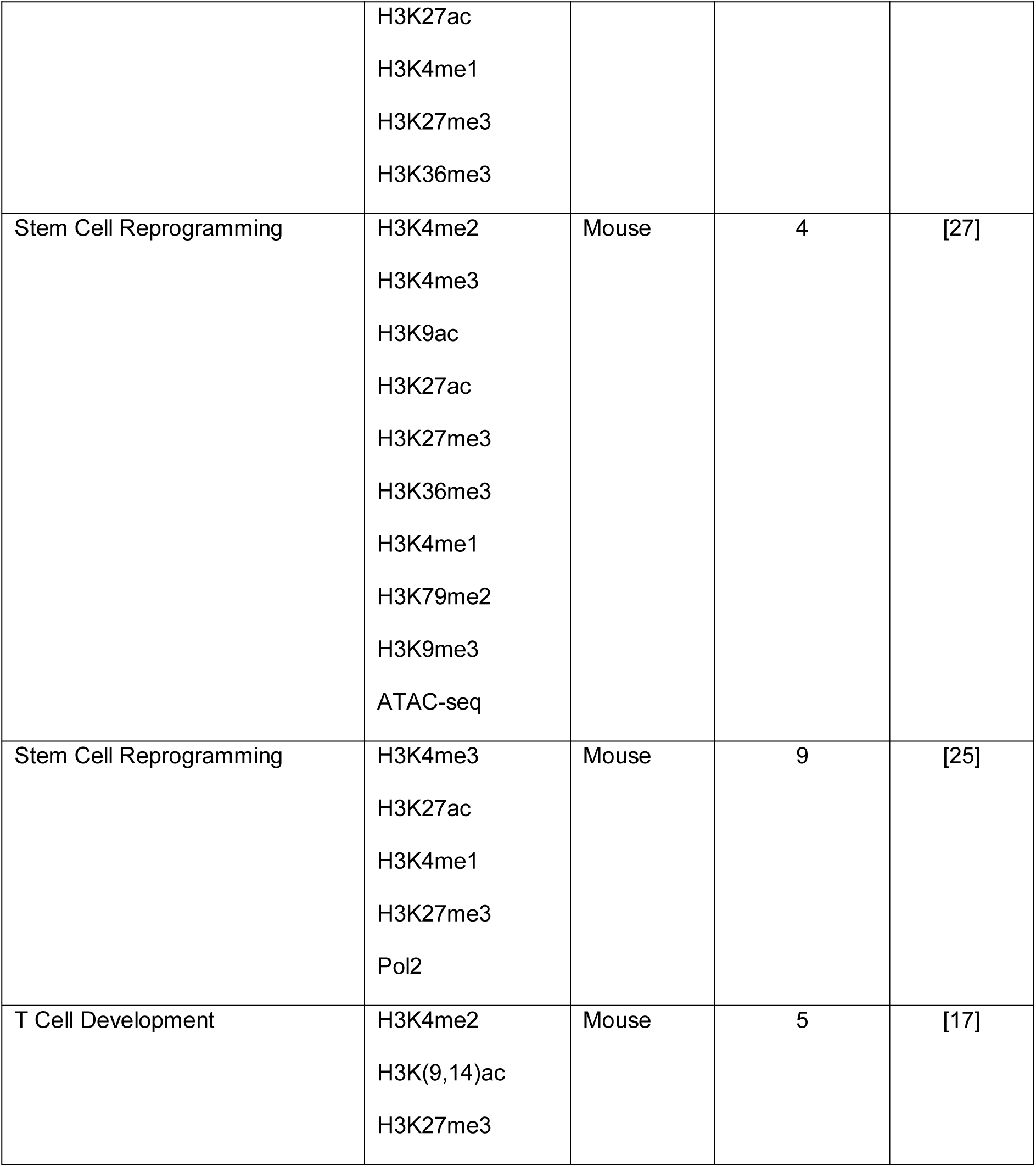
Datasets used for analysis with ChromTime

**Figure 7:**
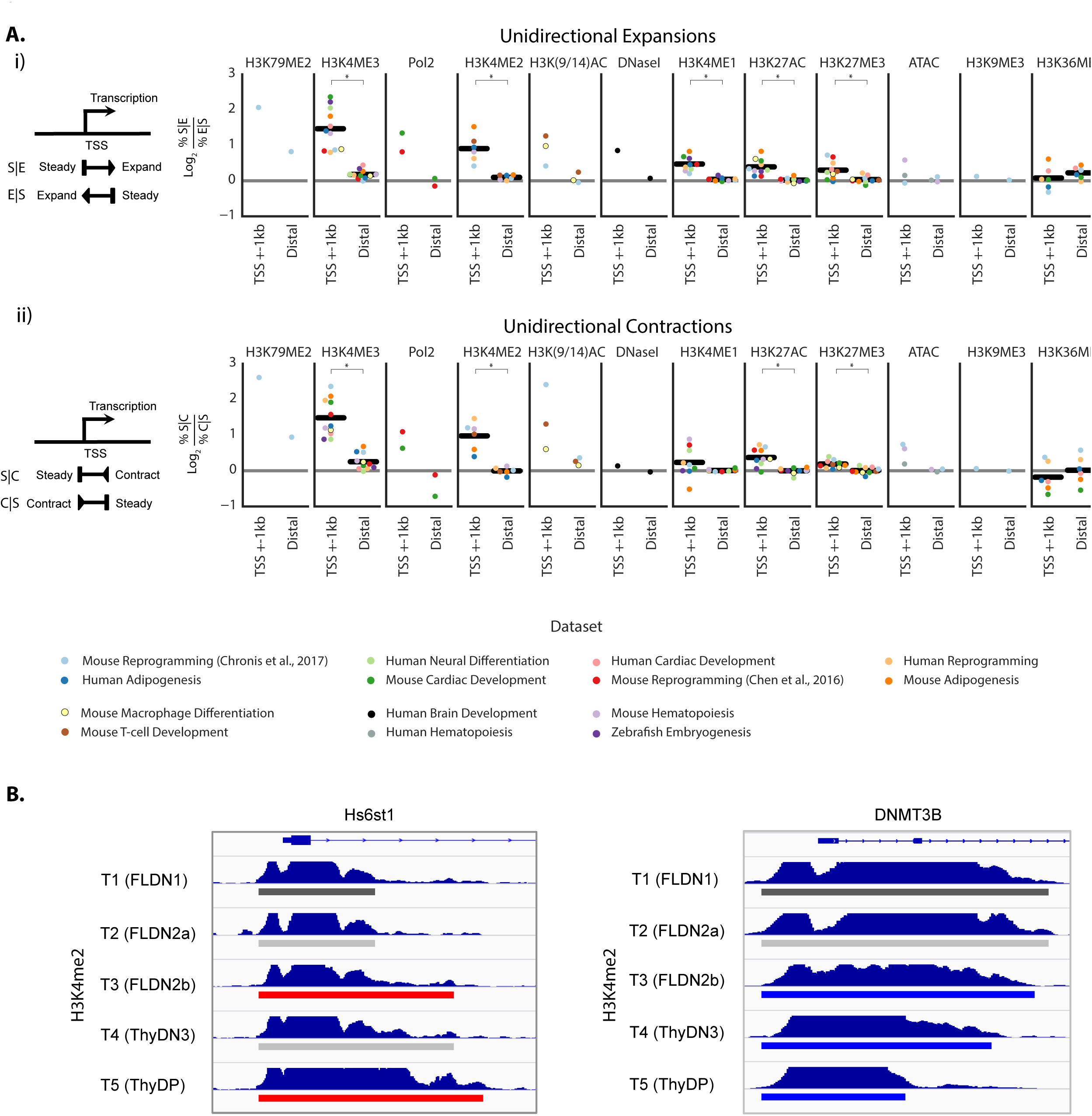
Direction of asymmetric dynamics correlates with direction of transcription. **(A) (i)** Left panel shows a schematic representation of unidirectional expansions that expand in the same direction as transcription and in the opposite direction of transcription. The adjacent plots show, for each mark, the average log_2_ ratio across all time points in each time course between the fraction of unidirectional expansions that expand in the same directions as transcription of the nearest gene and the fraction of unidirectional expansions that expand in the opposite direction of transcription of the nearest gene for blocks that are within 1 kb of annotated TSSs and for more distal blocks. Positive values correspond to enrichment of unidirectional expansions in the same direction as transcription. For marks mapped in at least six time courses, a black line is plotted representing the average across all data sets and significant differences are denoted with asterisks based on a two-tailed Mann-Whitney test at a p-value threshold of 0.05. **(ii)** Left panel shows analogous schematic for unidirectional contractions. Likewise, adjacent plots show, for each mark, the average log_2_ ratio between the fraction of unidirectional contractions that contract in the opposite direction of transcription of the nearest TSS and unidirectional contractions that contract in the same direction as transcription of the nearest TSS. **(B)** Left panel shows an example of unidirectional expansions between pairs of time points that expand in the same direction as transcription at the Hs6st1 gene of the H3K4me2 mark in the T cell development dataset[17]. Right panel shows an example of unidirectional contractions in the opposite direction of transcription at the DNMT3B gene. Time points 1, 2 and 3 correspond to in vitro differentiated T cell precursors, whereas time points 4 and 5 correspond to in vivo purified thymocytes. The predicted ChromTime peaks colored by their boundary dynamics are shown under the signal track for each time point.

## DISCUSSION

In this work, we presented ChromTime, a novel computational method for systematic detection of expanding, contracting and steady peaks of chromatin marks from time course high-throughput sequencing data. ChromTime employs a probabilistic graphical model that directly models changes in the genomic territory occupied by single chromatin marks over time. This approach allowed us to directly encode our modeling assumptions about dependencies between variables in an interpretable and extendable framework.

We applied ChromTime on ChIP-seq data for broad and narrow HMs and for Pol2, and on ATAC-seq and DNase-seq data from a variety of developmental, differentiation and reprogramming courses. Our results show that the method can identify sets of expanding and contracting peaks that are biologically relevant to the corresponding systems. In particular, expansions and contractions associate with up- and down-regulation of gene expression and differential TF binding, supporting the biological relevance of ChromTime predictions.

ChromTime gains power by both reasoning jointly about all time points in a time course and by explicitly modeling the peak boundary movements. Supporting this, in our analyses we observed that territorial changes identified by ChromTime had better agreement with gene expression changes compared to considering directly the boundary change of peaks called on data from individual time points in isolation. Additionally, we also observed for a range of cases that expanding and contracting peaks associated on average with greater change in gene expression compared to peaks with steady boundaries even after controlling for signal density changes. Some of the power that ChromTime gains from considering spatial information might be explained by its ability to differentiate territorial expansions or contractions which can reflect changes in the number sites of TF binding in close vicinity from changes in signal density within steady peak boundaries. Changes in signal density without territorial expansions or contractions might reflect a change in the proportion of cells with the chromatin mark without large changes in activity in any one cell. Additional power can come from the temporal and spatial information that allows the model to effectively smooth over noise in the data, thus enabling more biologically relevant inferences.

ChromTime enables novel analysis of directionality of spatial epigenetic dynamics. In this context, we found that asymmetric unidirectional expansions and contractions for several marks correlate strongly with direction of transcription in promoter proximal regions, which suggests that spatial dynamics at such locations may be related to actions of the transcriptional machinery. One possible explanation for the observed correlation between the direction of spatial dynamics of at least some HMs and transcription can be provided in part by previous studies that have shown that the Pol2 elongation machinery can recruit H3K4-methyltransferases such as members of the SET[55] and MLL[56] families at the promoters of genes. Our findings are consistent with such models where the Pol2 complex itself may be facilitating the attachment and removal of these marks[57].

The ChromTime software is also relatively efficient particularly when using its option to parallelize all computations during the parameter learning and prediction phases over multiple CPU cores. In our tests, processing ChIP-seq data for the H3K4me2 mark and control data from 5 time points in mouse T cell development[17] took 3 hours on a laptop computer using 4 CPU cores.

We applied ChromTime to a range of data types, however we found no single setting of the method options to be preferred in all cases (**Methods**). We thus created three modes with different default options: punctate mode used for ATAC-seq and DNase-seq, narrow mode used for ChIP-seq of narrow HMs, and broad mode used for ChIP-seq of broad HMs and Pol2. In principle, ChromTime can also be applied on ChIP-seq data of sequence specific TFs in punctate mode. However for this data, where binding can often be associated with a single point source such as individual instances of DNA sequence regulatory motifs, methods that predict the single point source across time points and the binding intensity associated with the source at each time point may be a more natural way to model the data.

Another limitation of the ChromTime method is that while the runtime of ChromTime still scales linearly with the number of time points, *T*, the number of observed combinations of dynamics can scale exponentially with *T*. This exponential growth can complicate downstream analyses that directly consider each combination of dynamics, as there will be *3^T-1^* possible sequences of dynamics at each side of a peak. Extensions of the ChromTime model could model the large number of combinations as being instances of a smaller number of more distinct dynamic patterns.

## CONCLUSIONS

The increasing availability of time course chromatin data provides an opportunity to understand chromatin dynamics in many biological systems. To facilitate reaching this goal we developed ChromTime, which systematically detects expanding, contracting, and steady peaks allowing extraction of additional information from this data. ChromTime gains power by both reasoning about data from all time points in the time course and by explicitly modeling movements of peak boundaries. We showed that ChromTime predictions associate with relevant genomic features such as changes in gene expression and TF binding. We demonstrated that territorial changes of peaks can contain additional information beyond signal density changes with respect to gene expression of proximal genes. ChromTime allows for novel analysis of directionality of spatial dynamics of chromatin marks. In this context, we showed for multiple chromatin marks that the direction of predicted asymmetric expansions and contractions of peaks strongly associates with direction of transcription in proximity of TSSs. ChromTime is generally applicable to modeling time courses of chromatin marks, and thus should be a useful tool to gaining insights into dynamics of epigenetic gene regulation in a range of biological systems.

## METHODS

### Overview of the ChromTime method

ChromTime takes as input a set of files in BED format with genomic coordinates of aligned sequencing reads from experiments for a single chromatin mark from a high-throughput sequencing experiment such as ChIP-seq, ATAC-seq or DNase-seq over a time course and, optionally, from a set of control experiments. ChromTime consists of two stages (**Fig 1B-C**):

1. Detecting genomic intervals (blocks) potentially containing regions of signal enrichment (peaks)
2. Learning a probabilistic mixture model for boundary dynamics of peaks within blocks throughout the time course and computing the most likely spatial dynamic and peak boundaries for each block throughout the whole time course

## Detecting genomic blocks containing regions of signal enrichment

The aim of this stage is to determine approximately the genomic coordinates of regions with potential peaks of signal enrichment at any time point in the time course (**Fig S1A-B**). The signal within these blocks will be used as input to build the mixture model in the next stage of ChromTime. ChromTime supports analysis of punctate, narrow and broad marks in three different modes, which are defined by different default options. The method partitions the genome into non-overlapping bins of predefined length, BIN_SIZE (by default, 200 bp in narrow and punctate modes, 500 bp in broad mode) and counts for each bin and time point the number of sequencing reads whose alignment starting positions after shifting by a predefined number of bases (SHIFT, 100 bp in the direction of alignment by default) are within its boundaries. Next, each bin at each time point is tested for enrichment based on a Poisson background distribution at a predefined false discovery rate (5% by default). The expected number of reads for a bin at position *p* and time point *t*, *λ*_*t,p*_, in the Poisson test is computed conservatively as the maximum of:

1. If control reads are provided: for each of windows of size *w=*1,000bp, 5,000bp and 20,000bp the average number of control reads in the window centered at the current bin, normalized by the ratio of total reads in the foreground and control experiments, that is:

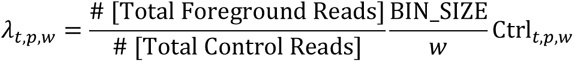

where Ctrl_*t,p,w*_ is the total number of control reads in each of window of size *w* around the bin at position *p* at time point *t*.
2. The average number of foreground reads per genomic bin.
3. 1 read.

Testing multiple different windows sizes for the background is a strategy we adopted from the MACS2 peak caller[38].

Within each time point, consecutive bins that are significantly enriched are merged into continuous intervals. The intervals are further extended in both directions to include continuous stretches of bins where each bin is significant based on a Poisson background distribution at a weaker P-value threshold (0.15 by default). Extended intervals within a predefined number of non-significant bins, MAX_GAP (3 bins by default), are further joined together. This joining strategy has been previously implemented by other peak callers for single datasets such as SICER[40]. Next, overlapping intervals across time points are grouped into blocks. To capture more of the potential background signal and to increase the likelihood that central bins within blocks contain higher foreground signal, the start and end positions of each block are extended additionally by a predefined number of bins, BLOCK_EXTEND, (5 by default) upstream from the left-most coordinate and downstream from the right-most coordinate of the intervals in the block, respectively, or up to the middle point between the current block and its adjacent blocks if they are within BLOCK_EXTEND bins apart. Restricting BLOCK_EXTEND to a relatively limited number of bins helps in keeping the running time of the method within reasonable bounds.

In narrow and punctate modes, blocks that contain multiple intervals at the same time point separated by gaps of non-significant bins longer than MAX_GAP are split into sub-blocks at each gap between those intervals. In particular, all gaps within a block are intersected across the time points that have gaps. For each gap intersection, the block is split at the position with the lowest average foreground signal across all time points. In broad mode, no such splitting is performed in order to avoid excessive peak fragmentation.

### Probabilistic mixture model for boundary dynamics of peaks within blocks across the time course

The foreground and the expected signal within the blocks are used as input to build a probabilistic mixture model for the boundary dynamics of the peaks within blocks (**Fig S1C**). One core assumption of the model is that each block contains at each time point exactly one peak, which can potentially have a length of zero bins. This implies that at each time point, the bins within a block can be partitioned into three continuous intervals: left-flanking background, foreground peak, and right-flanking background. For the bin in block *i*, at time point *t* and position *p*, let *O*_*i,t,p*_ denote the random variable that models the number of observed foreground reads, and let *O*_*i,t,p*_ denote the corresponding observed read counts. Let *V*_*i,t,p*_ denote the random variable for the label of the corresponding bin, which can either have the value PEAK or BACKGROUND. Let *X*_*i,t,p*_ denote a random variable for the vector of covariates for the corresponding bin, and *X*_*i,t,p*_ their corresponding values. The distribution of *O*_*i,t,p*_ conditioned on *V*_*i,t,p*_ and *X*_*i,t,p*_ is modeled with different negative binomial distributions depending on the value of *V*_*i,t,p*_ and *X*_*i,t,p*_:

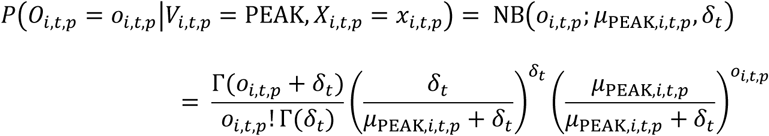

and

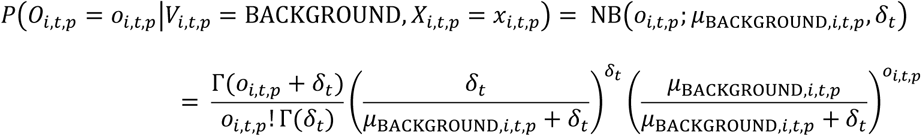

where *δ_t_* is the dispersion parameter. Similarly to negative binomial regression models[58], ChromTime models the mean of each component through the log link as a linear combination of a two-dimensional vector of covariates, *X*_*i,t,p*_ = (1, log *λ*_*i,t,p*_), which includes a constant term and the logarithm of the expected number of reads in the bin as computed in the previous section, *λ*_*i,t,p*_:

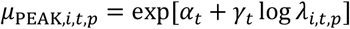

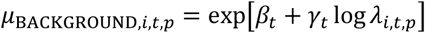

where *α_t_ β_t_* and *γ_t_* are time point specific scalar parameters. Negative binomial distributions have been successfully employed in a similar manner to capture the over-dispersion of sequencing reads in peak callers for single samples such as ZINBA[59]. Of note however, ChromTime requires that the dispersion parameter *δ_t_* and the coefficient *γ_t_* are shared between the two components at each time point. The first requirement ensures that the distribution with the smaller mean value has higher probabilities compared to the distribution with the larger mean value for the lowest values of the support domain of the negative binomial distribution, and that the opposite holds for the largest values of the support domain (**Supplementary Methods**). Sharing the dispersion parameter here is analogous to sharing the variance parameter in Gaussian mixture models. The second requirement to share the *γ_t_* parameter ensures that the control signal has equal importance in each component.

Formally, let *B_i,L,t_* and *B_i,R,t_* denote the random variables corresponding to the first and the last bin, respectively, in the peak partition at time *t* for block *i* relative to the beginning of the block, and let *N_i_* be the length of the block. We then have 1 ≤ *B_i,L,t_* ≤ *N_i_* + 1 and 0 ≤ *B_i,R,t_* ≤ *N_i_*, with values of *B_i,L,t_* = *N_i_* + 1 and *B_i,R,t_* = 0 corresponding to the special cases of starting a peak after all positions and ending a peak before all positions in a block, respectively. For *B_i,L,t_* and *B_i,R,t_* to denote valid interval boundaries, ChromTime also requires that *B_i,L,t_* ≤ *B_i,R,t_* + 1 at each time point. These constraints can be formally encoded by introducing one auxiliary binary variable for each time point in the model, *Z_i,t_*, such that:

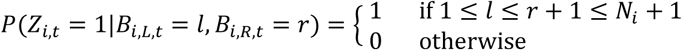

and thus also

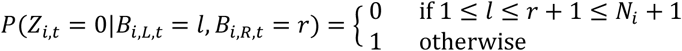

ChromTime treats all *Z_i,t_* variables as observed with values equal to 1 for all blocks and time points.

The conditional probability of the bin labels, *V_i,t,p_*, given the peak boundaries, *B_i,L,t_* and *B_i,R,t_*, are defined to be:

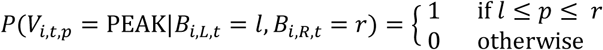

and thus also

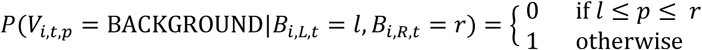

The probability of the observed read counts at time *t*, ***o**_i,t_* and *Z_i,t_* = 1, conditioned on the values of the peak boundaries, *B_i,L,t_* and *B_i,R,t_*, and the covariates at time point *t*, ***x**_i,t_*, under the model is then:

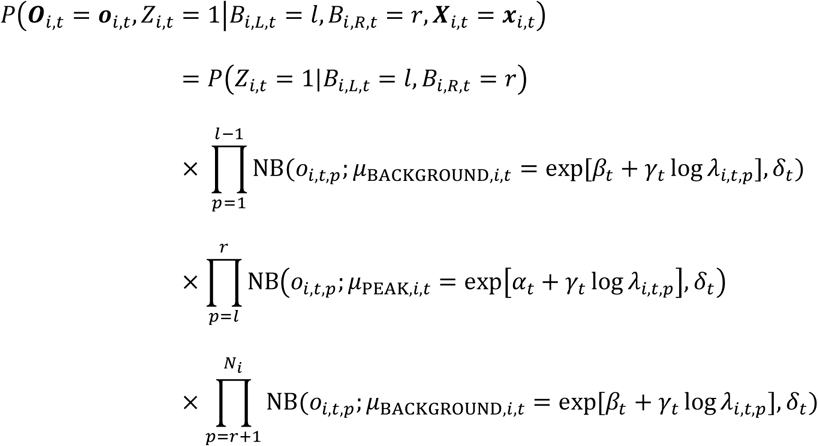

An important special case of the above formulation when *B_i,L,t_* = *B_i,R,t_* + 1 corresponds to modelling the whole signal at time point *t* as background, which enables ChromTime to accommodate time points that are all background by modelling them with zero length peaks. For this reason, ChromTime blocks internally have the same number of peak boundaries at all time points even if some time points are predicted as zero length peaks (i.e. all background). Boundaries of zero length peaks are treated by the model in the same way as boundaries of non-zero length peaks.

ChromTime assumes uniform prior probabilities for the left and the right end boundaries at the first time point:

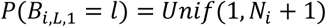

and

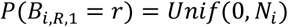

where *Unif(a, b)* denotes the uniform distribution of integer numbers in the closed interval [a, b].

Let *D_i,s,t_* denote the dynamic between time points *t* and *t+1* on boundary side *s*, where *s* is one of *L* (left side) or *R* (right side). Between any two time points the ChromTime model allows for one of three possible dynamics at both the left and the right end boundaries of a peak: STEADY, EXPAND or CONTRACT. To capture the change of boundary positions between consecutive time points *t* and *t+1* we define the quantities *J_i,L,t_* = *B_i,L,t_* − *B_i,L,t+1_* and *J_i,R,t_* = *B_i,R,t_* − *B_i,R,t+1_* corresponding to the left and right boundaries respectively. Positive values of *J_i,L,t_* and *J_i,R,t_* indicate the number of bases a peak expanded, whereas negative values indicate the number of bases a peak contracted, and a value of 0 indicates the peak held steady on the left and the right side respectively. ChromTime models *J_i,L,t_* and *J_i,R,t_* with different probability distributions for each of the three dynamics. For STEADY dynamic, ChromTime uses the Kronecker delta function:

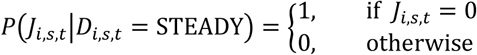

For expanding and contracting dynamics, ChromTime employs negative binomial distributions to model the number of genomic bins a peak boundary moves relative to the minimal movement of one bin required for peak expansions and contractions:

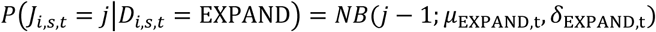

and

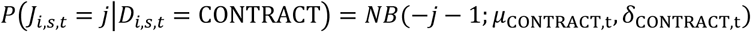

Furthermore, each distribution is parametrized with a mean and dispersion parameter depending on the dynamic and the time point, *t*: *μ*_EXPAND,t_, *δ*_EXPAND,t_ for expansions, and *μ*_CONTRACT,t_, *δ*_CONTRACT,t_ for contractions. Of note, in negative binomial distributions the probabilities for negative integers are defined to be 0. Therefore, the above parametrization enforces that boundary movements of negative or zero length (i.e. contracting or steady, respectively) are impossible for expansions and that boundary movements of positive or zero length (i.e. expanding or steady) are impossible for contractions.

The ChromTime model additionally assumes that there is a prior probability to observe each dynamic between time points *t* and *t+1*, *P*(*D_i,s,t_* = *d*) = *π_t,d_*, which is the same at each side (left and right). Users have the option to set a minimum prior probability (MIN_PRIOR) for the dynamics for all time points. This parameter can be used to avoid learning priors too close to zero, which in some cases can occur for more punctate marks where the short length of the peaks can cause the prior to become a dominant influence on the class assignment of the spatial dynamics. By default, MIN_PRIOR=0 in narrow and broad modes and MIN_PRIOR=0.05 in punctate mode.

For a time course with *T* time points we can express for block *i* the probability of a particular sequence of dynamics and boundary positions on the left side (***d**_L_* and ***b**_L_* respectively) and on the right side (***d**_R_* and ***b**_R_* respectively), and observing foreground counts ***o**_i_* and ***Z**_i_* = **1** conditioned on the values of the covariates, ***x**_i_* as:

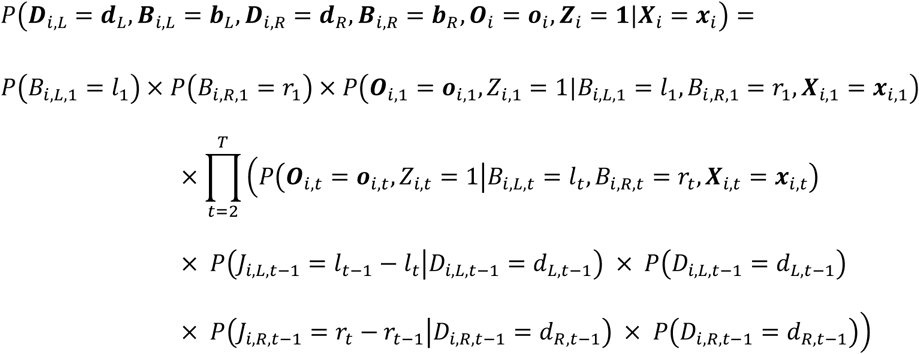

where ***Z**_i_* = **1** is used to denote *Z_i,t_* = 1 for all *t, d_s,t_* for t=1,…, *T-1* is the dynamic label for the *t*^th^ pair of consecutive time points on the left or the right side (*s = L* or *R*), respectively. Also ***b**_L_* and ***b**_R_* are the vectors of *T* boundary positions containing *l_t_* and *r_t_* for t=1,…, *T*, respectively.

The total probability of the signal in a block can be expressed as a sum over all possible sequences of dynamics and peak boundary positions that can generate the block across all time points. Thus, the probability of block *i* having observations ***o**_i_* and ***Z**_i_* = **1** given the covariates ***x**_i_*:

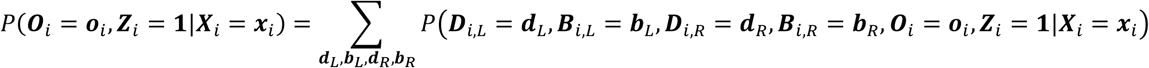

where ***d**_L_* and ***d**_R_* each iterate over all possible *3^T-1^* combinations of peak boundary dynamics, and ***b**_L_* and ***b**_R_* each iterate over all possible ways to place left and right end boundaries across all time points that are consistent with the requirements that 1 ≤ *B_i,L,t_* ≤ *B_i,R,t_* + 1 ≤ *N_i_* + 1 at each time point.

Let ***o*** be the total set of observed read counts in all blocks in the data, ***x*** be the set of the corresponding two-dimensional vectors containing the constant term and the log of the expected number of reads at each position and time point for each block, ***Z** = **1*** denotes all ***Z_i_*** = **1**, and *M* be the total number of blocks. Then, the likelihood of all blocks conditioned on their covariates is:

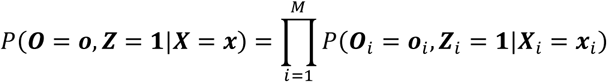

We note that in the above formulation allows ChromTime to model the appearance of a peak, if it occurs after the first time point in the time course, as an expansion from a zero length peak at the previous time point. Similarly, the disappearance of a peak is modelled as a contraction to a zero length peak at the next time point.

### Model optimization

The total set of parameters of the model consists of:

1. Prior probabilities of each dynamic *d* at each time point *t*: *π_t,d_*.
2. Parameters of the negative binomial distributions that model the PEAK and the BACKGROUND components at each time point: α_t_ β_t_ γ_t_ and δ_t_.
3. Parameters of the negative binomial distributions that model the boundary movements in EXPAND and CONTRACT dynamics at each time point: *μ*_EXPAND,t_, *δ*_EXPAND,t_ and *μ*_CONTRACT,t_, *δ*_CONTRACT,t_ respectively.

The optimal parameter values are attempted to be estimated by Expectation Maximization (EM). In particular, ChromTime attempts to optimize the conditional log-likelihood of the observed counts and ***Z**_i_* = **1** given the covariates (**Supplementary Methods**):

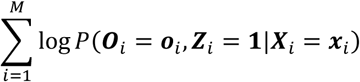

### Computing the most likely spatial dynamic and peak boundaries for each block across the whole time course

After the optimal values for all model parameters are estimated from the data, for each block the most likely positions of the peak boundaries at each time point are calculated. This procedure consists of two steps. First, ChromTime determines for each block all time points with significantly low probability of containing a false positive non-zero length peak. Second, conditioned on those time points, ChromTime computes the most likely assignment of the peak boundary variables at each side and each time point (**Supplementary Methods**).

### ChromTime options used in this study

In this work, we applied ChromTime in narrow mode on all data for H3K4me2, H3K4me3, H3K27ac and H3K(9,14)ac marks. We applied ChromTime in punctate mode on all ATAC-seq and DNase-seq data. No control reads were used for ATAC-seq and DNase-seq. In addition, foreground reads for ATAC-seq were shifted by 5bp in the direction of alignment (SHIFT=5), and for DNase-seq no shifting was applied (SHIFT=0). We applied ChromTime in broad mode on all data for H3K79me2, Pol2, H3K4me1, H3K27me3, H3K9me3 and H3K36me3 marks. All other options were set to their default values.

### Timing Evaluation

The timing evaluation was conducted on a MacBook Pro laptop with 2.7GHz Intel Core i7 and 16 GB RAM using 4 CPU cores.

### Analyses with External Data

The procedures for analyses with external are described in **Supplementary Methods**.

#### ABBREVIATIONS

ATAC-seq: Assay for transposase accessible chromatin coupled with high-throughput DNA sequencing
ChIP-seq: Chromatin immunoprecipitation coupled with high-throughput DNA sequencing
DHS: DNase I Hypersensitive Site
DNase-seq: DNase I hypersensitivity assay followed by high-throughput DNA sequencing
EM: Expectation Maximization
FDR: False Discovery Rate
HM: Histone mark
TF: Transcription factor
TSS: Transcription start site

## DECLARATIONS

### Authors’ contributions

J.E. conceived and supervised the project, designed the method, proposed analyses and wrote the manuscript. P.F. designed the method, implemented the software, proposed analyses, performed all analyses, processed all data and wrote the manuscript. Both authors read and approved the final manuscript.

### Ethics approval and consent to participate

Not applicable

### Consent for publication

Not applicable

### Availability of data and materials

ChromTime software is freely available at: https://github.com/ernstlab/ChromTime

No new experimental datasets were generated within this study. All datasets used as input for ChromTime are publicly available from the references listed in **Table 1**. Links to all other datasets (gene expression, TF binding and DHSs) used in this study are provided in the **Supplementary Methods**.

## Acknowledgements

We are grateful to Constantinos Chronis, Kathrin Plath and members of the Ernst lab for useful discussions.

